# A Nanopore-based method for generating complete coding region sequences of dengue virus in resource-limited settings

**DOI:** 10.1101/499111

**Authors:** Samuel Stubbs, Barbara Blacklaws, Benediktus Yohan, Frisalita A Yudhaputri, Brian Schwem, Edsel M Salvaña, Raul V Destura, Khin S Myint, R. Tedjo Sasmono, Simon D W Frost

## Abstract

Dengue virus (DENV) sequencing is a vital tool for surveillance and epidemiology studies. However, the current methods employed for sequencing DENV are expensive, laborious and technically demanding, often due to intra- and inter-serotype variability. Therefore, on-site DENV sequencing is not feasible in many of the areas where DENV is endemic. Surveillance in these areas can only be performed by shipping samples to well-equipped central laboratories for sequencing. However, long periods of inadequate storage and unreliable shipping conditions mean that such samples can arrive degraded, rendering sequence recovery difficult. We therefore aimed to develop an approach that is simple, portable and effective, to be used for on-site DENV sequencing in limited resource settings.

To achieve this, we first used the ‘Primal Scheme’ primer design tool to develop a simple and robust protocol for generating multiple short amplicons, covering the complete coding-region of DENV isolates. We then paired this method with the Nanopore MinION, a portable and affordable sequencing device, well-suited to minimal resource settings.

The multiplex PCR method produced full-coding-region coverage of all DENV samples tested with no optimisation required, and Nanopore sequencing of the short amplicons generated consensus sequences with high accuracy (99.52 - 99.92 %). Phylogenetic analysis of the consensus sequences generated using the new method showed that they formed monophyletic clusters with those produced by the current, long-amplicon, Illumina method, thus demonstrating that the two approaches are comparable. The multiplex method’s simplicity and portability compared to the current DENV sequencing approach make it well-suited for use in resource-limited, DENV-endemic regions. Deployment of the method in these regions would increase the capacity for DENV surveillance and has the potential to provide vital resolution for future DENV epidemiology studies.

## Introduction

Dengue virus (DENV) is a positive-sense RNA virus of the family *Flaviviridae*. It is a mosquito-borne pathogen which can predominantly be found circulating in urban or semi-urban areas in tropical regions, where its vectors, *Aedes aegypti* and *Aedes albopictus*, are present (1). DENV has been estimated to infect 390 million people each year, approximately 100 million of whom exhibit clinical symptoms (2). However, the true global burden of dengue has been difficult to ascertain due to limited surveillance and diagnostic infrastructure in endemic regions across much of Asia, South America, the Pacific and the Caribbean (3).

There are four antigenically distinct serotypes of DENV1-4 which share approximately 50-70 % homology at the nucleotide level (4). In many dengue-endemic regions, all four serotypes co-circulate (5). Infection by DENV elicits life-long immunity against the infecting serotype, but does not confer lasting protective immunity against the remaining three serotypes. Dengue outbreaks in endemic regions of the Americas and South East Asia are seasonal, with larger epidemics occurring every 3-5 years (3,6). Several of these epidemics have been associated with switches in the dominant circulating clade (7–9), suggesting that dengue virus surveillance has the potential to be a useful tool for the prediction and early identification of outbreaks.

DENV genome sequencing can be used to provide insights into how the virus is being transmitted and how it is spread (10). An understanding of these factors can inform public health policies by identifying the source of an outbreak or regions at high risk of outbreaks in the future. These regions can then be targeted with additional vector control or surveillance measures. Furthermore, if viral outbreaks can be identified early, it may enable these measures to be applied quickly enough to prevent significant spread. Monitoring the evolution of DENV is also important, as amino acid differences between closely related genotypes have been shown to influence both the virulence and transmissibility of the virus (11).

For a sequencing method to be rapid (i.e. a turnaround of days rather than weeks), slow and laborious viral culture methods must be avoided. Direct sequencing of DENV from clinical samples is possible, however, unless the viral copy number is high, this approach will predominantly produce sequences derived from the host and very few from the infecting virus. If sequence coverage of the virus is insufficient, the viral genome cannot be accurately reconstructed, and phylogenetic analysis becomes unreliable. One method used to overcome this lack of sensitivity is to use (RT-)PCR to specifically amplify the viral genome prior to sequencing. Whole genome sequencing of DENV is typically performed in this way, using serotype-specific primer sets to produce 5 amplicons, approximately 2 - 2.5 kb in length, which overlap to cover the entire genome (12,13). The amplicons from these 5 single-plex reactions are typically visualised by gel electrophoresis, purified, and prepared for sequencing on the Illumina platform.

However, a lack of resources means that the current single-plex PCR and Illumina-sequencing approach is not viable in many of regions of Asia, Africa and Latin America where DENV is present, limiting viral surveillance efforts. The cost and expertise required for Illumina sequencing is currently restricted to modern, well-funded laboratories, whilst performing multiple single-plex reactions for every sample can be time consuming and cumbersome, particularly in under-equipped laboratories or in the field. Single-plex reactions are also susceptible to contamination, handling and pipetting errors, and costlier than multiplexing in terms of reagents. Furthermore, the production of long PCR amplicons requires the viral RNA genome to be largely intact. This makes the approach inappropriate for samples with low viral loads, as the likelihood of long fragments being present is reduced with decreasing concentration. Additionally, in many remote dengue endemic regions sample storage options are limited to −20 ° C freezers. Long-term storage of RNA in these conditions is likely to lead to sample degradation, rendering amplification impossible. In summary, on-site viral surveillance is not possible in many DENV endemic areas. Surveillance studies, when undertaken in these areas, must instead rely on samples being collected and transported to central sequencing centres under costly and unpredictable storage conditions.

One way to overcome the issue is to set-up further sites with sequencing facilities. However, many of the most commonly used next generation sequencing platforms such as Illumina, IonTorrent and PacBio, are prohibitively expensive to purchase and maintain and require a full-time, skilled technician to run the machines. The Oxford Nanopore Technologies MinION sequencing device is an affordable and highly portable alternative, which has the potential to be used for viral genomic surveillance at sites lacking the resources classically required for such work. Compared to the aforementioned platforms it requires relatively little infrastructure to set-up and run. An additional benefit of the Nanopore platform is that data in generated in real-time which allows rapid identification of the virus in an outbreak situation. The method could also be paired with novel, field-deployable, detection methods such as a CRISPR-cas9 approach (14), potentially well suited to harsh environments where the equipment necessary for DENV diagnosis or genotyping by conventional methods such as qPCR is not available.

Here we aimed to develop a method capable of rapidly generating complete coding sequences from DENV samples, which is suitable for resource-limited settings where DENV is prevalent. The method makes use of the ‘Primal Scheme’ multiplex-PCR approach developed for field sequencing of two other arboviruses: Zika virus and chikungunya virus (15). We tested the method on reference material as well as a group of clinical samples, both containing representatives of each of the four DENV serotypes. The resulting amplicons were sequenced in parallel on both the Nanopore MinION and the Illumina MiSeq, and consensus sequences generated using multiple read mapping programs. The resulting genome assemblies were compared to assess the approach in terms of viral genome coverage and assembly accuracy.

## Methods

### Design of multiplex primer sets

The ‘Primal Scheme’ primer design tool (primal.zibraproject.org) was used to design multiplex primer sets. These primer sets were designed to produce multiple 400 bp amplicons covering the entire coding region of DENV, with a 50 bp overlap between adjacent amplicons to facilitate assembly of the sequence data. Primer design was based on full genome sequences of isolates from Indonesia obtained from NCBI GenBank (www.ncbi.nlm.nih.gov/genbank) (a full list of primer sequences is given in the supplementary material). Multiple sequence alignment using Clustal Omega (16) revealed 68-73 % sequence similarity between serotypes, and 93-100 % similarity within serotypes within the database. Greater than 10 % nucleotide divergence between serotypes meant that it was necessary to design individual primer sets for DENV serotypes 1-4. Finally, sequences with > 99 % homology were removed to avoid primer selection bias, and the sequence of the most recent isolate was placed at the start of each FASTA file to act as the ‘most representative’ genome.

### Samples

The amplification and sequencing methods were first tested using the AmpliRun dengue RNA control set (VirCell, Granada, Spain), consisting of purified RNA from each of the four DENV serotypes. According to the manufacturer, these control samples ranged in concentration from 12,500 – 20,000 viral genome copies / µl. Their complete genome sequences are available in NCBI GenBank: DENV1 (KM204119.1), DENV2 (KM204118.1), DENV3 (KU050695.1), DENV4 (KR011349.2).

Four plasma samples, each representing one of the four DENV serotypes, were next used to test the efficacy of method on clinical samples. These samples were taken from patients in the Philippines in whom DENV infection had previously been confirmed and were obtained with informed, written consent under the approval of University of the Philippines Manila Research Ethics Board.

### RNA extraction and reverse transcription

RNA was extracted from 200 µl of each plasma sample using the High-Pure Viral Nucleic Acid kit (Roche) and eluted in 20 µl of elution buffer. 7 µl of extracted RNA were reverse transcribed using 50 ng of random hexamers (Invitrogen) and Superscript III enzyme (Thermo Fisher Scientific), as per the manufacturer’s instructions. Second-strand synthesis of cDNA was performed by adding 2·5 U (0.5 µL) of large Klenow fragment (New England Biolabs, NEB) to each reaction and incubating at 37 ° C for 60 min followed by 75 ° C for 10 min to inactivate the enzyme.

### PCR Amplification

Single-plex amplicons (1 - 2.5 kb in length) were produced using Q5 Polymerase (NEB) and serotype-specific primers following the protocols described by Ong et al (17) (DENV1-3) and Sasmono et al (18) (DENV4). Five amplicons were produced per sample in 5 separate 20 µL reactions, with 2 µL from the cDNA step used as direct input for each. 15 µL of each PCR reaction were visualised on a 1 % agarose gel to confirm successful amplification and repeated if unsuccessful. Once amplified, bands of the appropriate size were excised from the agarose gel and purified using the Monarch Gel Purification Kit (NEB), eluting in 10 µL of elution buffer.

Multiplex amplicons were produced in two reactions per sample, with the primers separated into two pools to avoid interference between overlapping amplicons. PCR amplification was performed using Q5 polymerase in a 25 µL reaction volume following the protocol described by Quick et al (15). The final individual primer concentration was 0.015 µM, and 2.5 µL of cDNA / reaction was used as input. 5 µL of each PCR reaction were visualised on a 2 % agarose gel to confirm successful amplification. Amplicons were purified from the remaining 15 µL PCR reaction by adding a 1 x volume of KAPA Pure beads (Roche), washing twice in 75 % ethanol and eluting in 30 µL of EB buffer (Qiagen).

### Nanopore sequencing and consensus sequence generation

Barcoded, 1D sequencing libraries were prepared using the Oxford Nanopore Native Barcode kit (EXP-NBD103) and the Oxford Nanopore 1D Ligation Sequencing kit (SQK-LSK108) following the protocol described by Quick et al (15).

The R9 flow-cell was QC-ed and primed following Oxford Nanopore’s 1D native barcoding protocol (available at https://community.nanoporetech.com/protocols). 75 µL of library mix consisting of 14.5 µL of barcoded library (∼ 50 - 75 ng), 35 µL of RBF, and 22.5 µL of LLB (both included in the 1D Sequencing Kit, Oxford Nanopore), was loaded onto the flow-cell and sequenced using the MinKNOW software v1.11.5 (Oxford Nanopore).

Raw FAST5 files were base-called and de-multiplexed using Albacore v 2.1.10 (Oxford Nanopore) with the settings: --flowcell FLO-MIN107 --kit SQK-LSK108 --barcoding -q 7. Porechop v 0.2.3 (available at: github.com/rrwick/Porechop) was used with default settings to trim sequencing adapters and perform more stringent demultiplexing. Serotype-specific primer sequences were trimmed from the ends of the de-multiplexed reads in both the forward and reverse orientation using Cutadapt v 1.16 (19) with the anchored adapter setting and a permitted error rate of 0.2. The de-multiplexed and trimmed FASTQ files were aligned to the appropriate dengue reference sequence (KM204119.1 for DENV1, KM204118.1 for DENV2, KU050695.1 for DENV3, and KR011349.2 for DENV4) using graphmap align v0.5.2 (20), BWA mem v0.7.17 (21), and minimap2 v2.9 (22).

The resulting BAM alignment files were sorted and indexed using samtools v1.7 (23). BAM alignment files were then imported into the CLC Genomics Workbench v7.5.1 (Qiagen) and the consensus sequences were extracted in FASTA format for analysis. Consensus sequences were called from all regions with at least 1 x coverage, and a threshold of 0.3 was set for calling ambiguities.

### Illumina library sequencing and consensus sequence generation

Illumina multiplex libraries were prepared from the purified amplicons using the NEBNext Ultra II kit and sequenced on the Illumina MiSeq using the v2 chemistry producing 2 x 250 bp paired-end reads.

Serotype-specific primer sequences were trimmed from the ends of the de-multiplexed reads in both the forward and reverse orientation using Cutadapt v1.16 with the anchored adapter setting, a permitted error rate of 0.1, and a minimum q-score of 30. Trimmed, paired-end read files were imported into CLC Genomics Workbench v7.5.1 for analysis. Reads were mapped to a reference sequence of the appropriate serotype requiring a minimum of 64 % identity (80 % similarity over 80 % of the read length). The consensus sequence was extracted from regions of the alignment with a minimum of 1 x coverage, and a threshold of 0.2 for calling ambiguities.

### Assessment of read and consensus sequence accuracy

The individual Nanopore read error rate was calculated by aligning the de-multiplexed, primer-trimmed reads to the corresponding Illumina, single-plex-generated consensus sequence using graphmap align v0.5.2. The consensus generated using the single-plex, Illumina sequencing method was employed as a reference sequence as it is the current standard, and is produced using high-accuracy reads with an even depth of coverage. The information contained in the ‘MD field’ of the resulting BAM alignment file was then parsed to count the number of mismatches and deletions. The resulting output was plotted in RStudio v1.1.383 (available at: www.rstudio.com) using R v3.4.3 (24), and the ggplot2 (25) and ggforce packages (available at: github.com/thomasp85/ggforce).

Accuracy of the multiplex and MinION consensus sequences was assessed in much the same manner as individual read accuracy. The consensus sequences were first divided into 100 bp fragments using the ‘chop_up_assembly’ script (available at: github.com/rrwick/Basecalling-comparison/blob/master/chop_up_assembly.py) so that accuracy could be assessed separately across each region of the genome. The 100 bp sequences were then aligned to their Illumina-generated equivalent and the ‘MD field’ of the resulting BAM alignment file was parsed to count the number of mismatches and deletions. The alignment was also checked by eye in order to correct for ambiguous base matches present in both the consensus and reference sequences, as these were seen as mismatches by the MD field parser.

### Phylogenetic analysis

Consensus sequence FASTA files were trimmed to the location of the most 3’ and 5’ multiplex primer binding-regions, and combined with reference sequences from their respective serotypes. Reference sequences were obtained from the Dengue Virus Typing Tool (available at www.krisp.org.za/tools.php). Multiple sequence alignments of the complete DENV coding region were generated using the FFT-NS-i method of MAFFT v7.407 (26). Phylogenetic trees were reconstructed using maximum likelihood using IQTREE v1.6.8 (27), using model selection (‘-m TESTMODELNEW’) (28) and a more thorough nearest neighbour interchange search (‘-allnni’). The resulting trees were visualised using ggtree (29) and distances between taxa were calculated using the cophenetic.phylo function in R.

### Data sharing

Raw sequence data have been deposited in the NCBI Sequence Read Archive database under accession number SRPXXXXXX [*accession number will be inserted in the R1 of the manuscript*].

## Results

### PCR amplification of DENV

An aliquot of each multiplex PCR reaction was run on an agarose gel to assess whether amplification had been successful. Visualisation revealed a 400 bp band had been generated for all 8 samples (4 control RNA samples and 4 clinical plasma samples) on the first attempt. For some samples, a larger band, approximately 1 kb in length, was also visible (Supplementary Figure 1). Bands of this size suggest that one or more primers were unable to anneal to the template; yet amplification of the region still occurred through use of a primer from a neighbouring region.

The single-plex approach consisted of five separate PCR reactions, meaning each amplicon could be checked individually. For the four control RNA samples, all 5 amplicons were produced, with no need for optimisation of the reaction or thermal cycling conditions. However, for several of the clinical samples, amplification of the 2 kb amplicons required adjustments to the reaction conditions. Indeed, amplification of the 3’ region of DENV1, 2 and 4 was only achieved by replacing the antisense primer with primers from the multiplex primer-sets.

### Illumina sequencing of multiplex and single-plex amplicons

The efficacy of the multiplex approach was next assessed by sequencing the amplicons on the Illumina MiSeq, generating between 606,126 – 933,018 250 bp PE reads per sample. The resulting reads were trimmed, and assembled by mapping to a serotype-specific reference. DENV genome coverage of between 96.12 – 100.00 % was achieved, with a mean depth of over 10,000 x for all samples (Table 1).

**Table 1.** Illumina sequencing results for DENV using the multiplex PCR (400 bp amplicon) approach. Consensus sequences assembled from the 400 bp data were compared to consensus sequences generated using the current standard single-plex PCR (2 kb amplicon) approach, in order to assess their accuracy.

| Sample Type | Genotype<sup>a</sup> | Reads<sup>b</sup> | Coverage (%)<sup>c</sup> | Depth<sup>d</sup> | Alignment Accuracy (%)<sup>e</sup> | Overall Accuracy (%)<sup>f</sup> |
| --- | --- | --- | --- | --- | --- | --- |
| Control | DENV1 - I | 933,018 | 100 | 15,249 | 99.98 | 99.98 |
| Control | DENV2 - IV | 786,724 | 100 | 13,331 | 99.84 | 99.84 |
| Control | DENV3 - V | 834,590 | 98.63 | 14,102 | 99.97 | 98.60 |
| Control | DENV4 - I | 606,126 | 96.74 | 10,437 | 99.89 | 96.63 |
| Clinical | DENV1 - IV | 660,900 | 99.4 | 11,166 | 99.97 | 99.37<sup>g</sup> |
| Clinical | DENV2 - II | 893,504 | 100 | 14,683 | 99.98 | 99.98<sup>g</sup> |
| Clinical | DENV3 - I | 798,382 | 100 | 13,283 | 99.97 | 99.97 |
| Clinical | DENV4 - II | 817,756 | 96.12 | 14,328 | 99.72 | 95.85<sup>g</sup> |

<sup>a</sup> Genotype assigned by phylogenetic analysis of the consensus sequence
<sup>b</sup> Number of 400 bp amplicon, Illumina reads generated for the sample
<sup>c</sup> Percentage of the dengue virus genome covered by at least one read
<sup>d</sup> Mean depth of coverage for the dengue virus genome
<sup>e</sup> Percentage of bases with > 1 x coverage that matched between the multiplex- and single-plex-generated consensus sequences when the two were aligned (regions with no coverage excluded)
<sup>f</sup> Overall similarity between the multiplex- and single-plex-generated consensus sequences (regions with no coverage counted as mismatches)
<sup>g</sup> The 3' end of the dengue virus genome could not be amplified for several of the clinical samples using the single-plex (2 kb amplicon) approach (see Coverage column). The consensus sequence for these samples was therefore truncated, and the accuracy of the multiplex consensus sequence for these missing regions could not be assessed.

In order to assess the accuracy of the multiplex approach, consensus sequences were aligned against those generated by the current single-plex approach. It should be noted that the consensus sequences for three clinical samples were incomplete due to failure of the single-plex primers to amplify the 3’ region (as described above). This meant that the accuracy of the multiplex consensus sequences could not be fully assessed for these regions. Considering regions with a minimum of 1 x coverage, the multiplex consensus sequences reached 99.72 – 99.98 % similarity to their single-plex counterparts. If lack of coverage was also considered an ‘inaccuracy’, 6/8 samples generated consensus sequences with 98.6 % similarity or higher. The two multiplex consensus sequences with less than 98% similarity to their single-plex counterparts were produced by the DENV4 control and clinical isolate (96.63 % and 95.85 %). These samples were also those with the largest regions of 0 x coverage (3.26 % and 3.88 %).

### Nanopore-sequencing of amplicons generated from DENV control RNA

Amplicons generated from the four DENV control samples using the multiplex and single-plex approach were next sequenced on the Nanopore MinION, in order to test the performance of the device in comparison to the Illumina MiSeq. The amplicons were barcoded for multiplex sequencing, and MinION sequencing was performed for 14 h, producing a total of 236,911 reads post-QC (7,271 – 57,562 reads per sample) (Table 2).

**Table 2.** Nanopore sequencing results for DENV control RNA samples, using both the single-plex PCR (2 kb amplicon) and multiplex PCR (400 bp amplicon) approaches.

| **Genotype<sup>a</sup>** | **Amplicon Size** | **Reads<sup>b</sup>** | **Genome Coverage (%)<sup>c</sup>** | **Average Depth (x)<sup>d</sup>** | **Alignment Length (bp)<sup>e</sup>** | **Consensus Accuracy (%)<sup>f</sup>** |
| --- | --- | --- | --- | --- | --- | --- |
| DENV1 - I | 2000 | 53,017 | 100 | 4452 | 10,736 | 99.99 |
| DENV2 - IV | 2000 | 7,271 | 100 | 929 | 10,723 | 99.79 |
| DENV3 - V | 2000 | 24,842 | 100 | 2207 | 10,696 | 100.00 |
| DENV4 - I | 2000 | 57,562 | 100 | 4145 | 10,644 | 99.98 |
| DENV1 - I | 400 | 18,783 | 100 | 444 | 10,736 | 99.92 |
| DENV2 - IV | 400 | 16,663 | 100 | 617 | 10,723 | 99.67 |
| DENV3 - V | 400 | 18,177 | 100 | 332 | 10,696 | 99.86 |
| DENV4 - I | 400 | 40,596 | 100 | 758 | 10,644 | 99.52 |

<sup>a</sup> Genotype assigned by phylogenetic analysis of the consensus sequence
<sup>b</sup> Number of Nanopore reads generated for the sample (post-QC)
<sup>c</sup> Percentage of the dengue virus genome covered by at least one read
<sup>d</sup> Mean depth of coverage for the dengue virus genome
<sup>e</sup> Length of sequence that could be aligned to the Illumina-generated, 2 kb consensus
<sup>f</sup> Similarity between the multiplex (400 bp amplicon) and singleplex (2 kb amplicon)-generated consensus sequences

Accuracy of the raw Nanopore reads was determined by aligning them to a reference sequence, generated using the conventional, single-plex PCR and Illumina sequencing approach. Mean accuracy of the Nanopore reads was comparable across all DENV serotypes and amplicon sizes (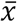 = 90.12 – 90.87 % for multiplex amplicons, 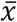 = 90.00 – 90.90 % for single-plex amplicons) (Figure 1A). However, there was greater variability in read accuracy in the data generated by the multiplex amplicons (stdev = 3.57 – 3.81 %) compared to the data generated by the single-plex amplicons (stdev = 2.24 – 3.07 %).

**Figure 1.**
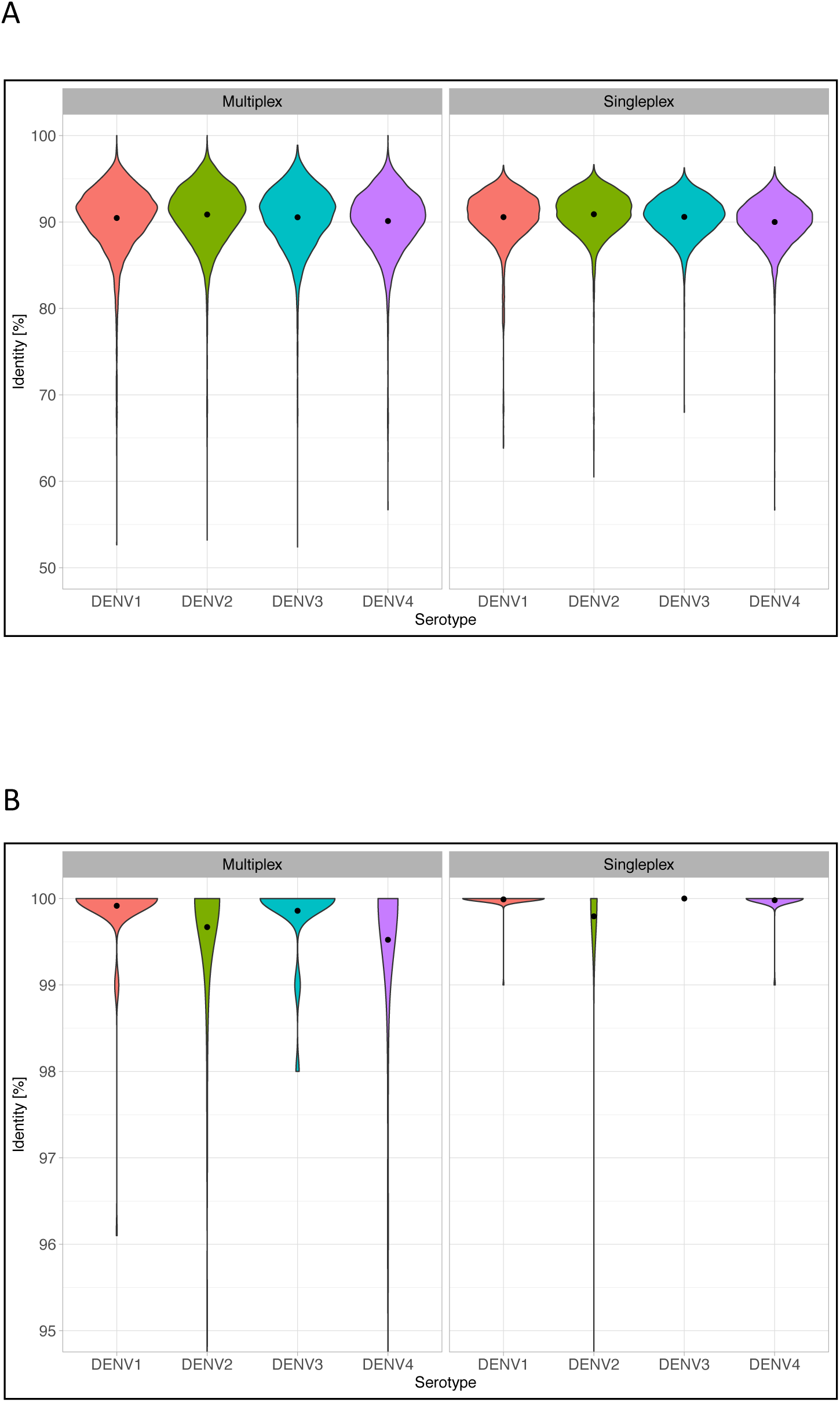
Nanopore sequencing read and consensus sequence accuracy. DENV RNA control samples were prepared using the multiplex PCR (400 bp amplicon) and single-plex PCR (2 kb amplicon) approaches, and sequenced on the Nanopore MinION. Errors such as SNPs, insertions, and deletions were identified by aligning the Nanopore-generated sequences to a reference sequence, which was generated using the current standard approach: Illumina-sequencing of single-plex (2 kb) amplicons. Percentage similarity to the Illumina reference is plotted on the y-axis. A) Accuracy of the individual Nanopore reads. Mean read accuracy (•) was approximately 90 %, and was comparable between the two approaches. B) Accuracy of Nanopore-generated consensus sequences. For all samples, assembly of the reads into a consensus resulted in a sequence more than 99.5 % identical to the Illumina reference (Mean consensus accuracy is indicated by the black dot (•)). The consensus sequences generated using the multiplex PCR approach were marginally lower accuracy than those generated using the single-plex approach.

Primer sequences and low-quality bases were trimmed from the ends of the Nanopore reads and the trimmed reads were assembled by mapping to a serotype-specific reference genome. Full coverage of the coding region was generated by both the multiplex and single-plex approaches for all four DENV serotypes. Mean coverage depth ranged between 332 x (DENV3) to 758 x (DENV4) for the multiplex amplicons, and between 929 x (DENV2) to 4452 x (DENV1) for the single-plex amplicons.

Consensus sequences were called from the alignments, divided into 100 bp fragments, and re-aligned to their intact, single-plex PCR, Illumina-generated, counterpart in order to determine their accuracy across the genome. All assemblies reached a mean accuracy of 99.5 % or more (Figure 1B) and the majority of 100 bp regions were 100 % accurate, with the majority of inaccuracies limited to a small number of regions. However, mean accuracy was on average 0.2 % lower for consensus sequences generated from the multiplex amplicons compared to their single-plex counterparts.

Visualisation of consensus accuracy across the DENV genome revealed that drops in accuracy tended to coincide with regions of low coverage depth, most noticeably when depth was less than 50 x (Figure 2). These areas of low coverage depth were also found to coincide with regions containing higher numbers of ambiguous bases in the derived consensus sequence (Supplementary Figure 2). Conversely, there were no regions of low coverage and fewer regions of low accuracy in the data-set generated using the single-plex approach (Supplementary Figure 3).

**Figure 2.**
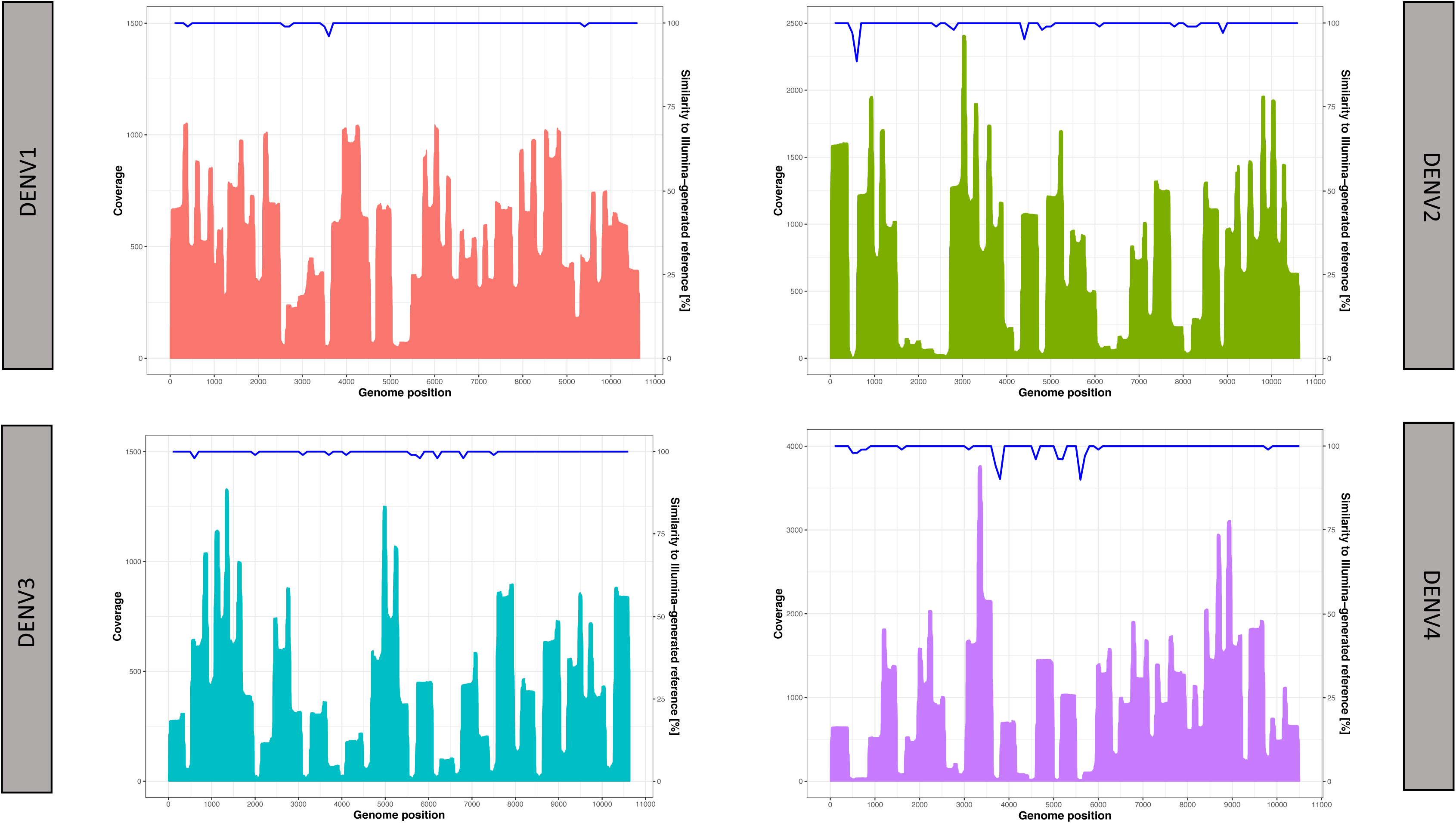
Nanopore sequencing coverage and accuracy for DENV control RNA samples, using the multiplex (400 bp amplicon) approach. Regions of reduced accuracy in the Nanopore-generated consensus sequence (indicated by the blue line and plotted on the right y-axis) were correlated with regions of low sequencing coverage (indicated by the area plot and plotted against the left y-axis). Consensus accuracies were determined by alignment against a reference sequence, generated using the standard, Illumina and single-plex (2 kb amplicon), approach.

### Nanopore sequencing of multiplex amplicons generated from clinical samples

Nanopore sequencing was next performed on amplicons generated from four DENV-infected plasma samples using the multiplex approach. The amplicons were barcoded, and multiplex sequencing was performed for 9 h, producing between 5,086 – 13,610 reads per barcode post-QC (Table 3).

**Table 3.** Nanopore sequencing results for DENV-infected clinical samples, using the multiplex (400 bp amplicon) approach.

| **Genotype<sup>a</sup>** | **Reads <sup>b</sup>** | **Genome**<br>**Coverage (%) <sup>c</sup>** | **Average**<br>**Depth (x) <sup>d</sup>** | **Alignment**<br>**Length (bp) <sup>e</sup>** | **Consensus**<br>**Accuracy (%) <sup>f</sup>** |
| --- | --- | --- | --- | --- | --- |
| DENV1 - IV | 5,087 | 100 | 205 | 10,398 | 99.09 |
| DENV2 - II | 21,859 | 100 | 792 | 10,395 | 99.51 |
| DENV3 - I | 7,974 | 100 | 307 | 10,655 | 99.41 |
| DENV4 - II | 13,610 | 100 | 486 | 9,706 | 98.52 |

<sup>a</sup> Genogroup assigned by phylogenetic analysis of the consensus sequence
<sup>b</sup> Number of Nanopore reads generated for the sample (post-QC)
<sup>c</sup> Percentage of the dengue virus genome covered by at least one read
<sup>d</sup> Mean depth of coverage for the dengue virus genome
<sup>e</sup> Length of sequence that could be aligned to the Illumina-generated, 2kb consensus (less than full length for DENV1, 2 and 4)
<sup>f</sup> Similarity between the Nanopore-generated consensus sequence using the multiplex approach, and a reference generated by Illumina sequencing using the single-plex approach

Reference-based assembly of the Nanopore reads generated 100 % coverage of the coding-region for all four DENV serotypes. Depth of coverage ranged from an average of 205 x (DENV1) to 792 x (DENV4). The accuracy of the Nanopore-generated consensus sequences was assessed by aligning them to consensus sequences generated using the Illumina single-plex approach. Mean consensus accuracy ranged from 98.52 – 99.51 %. Regions of reduced accuracy again coincided with regions of lower coverage, as previously observed in the control samples (Figure 3).

**Figure 3.**
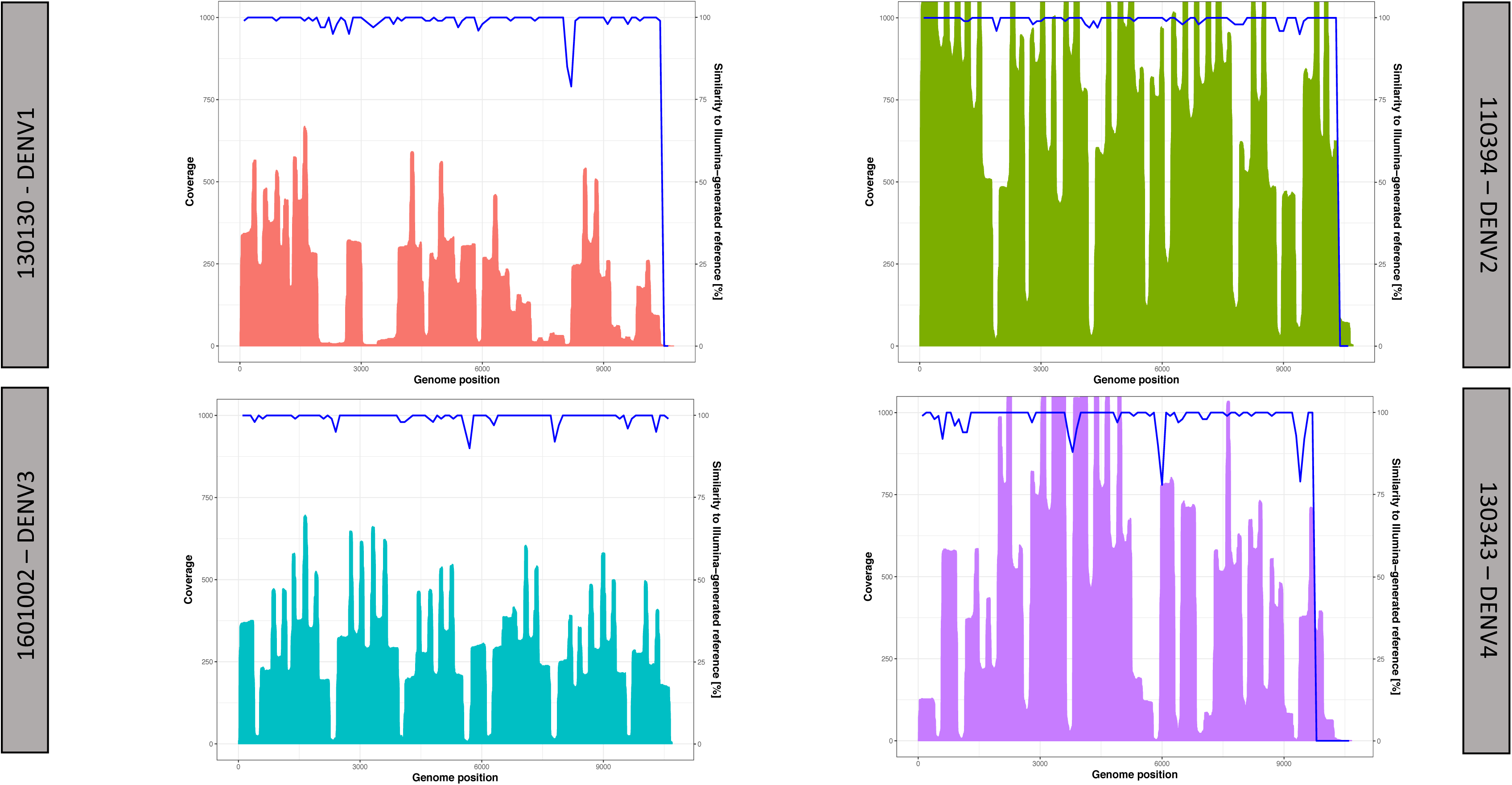
Nanopore sequencing coverage and accuracy for DENV clinical isolates, using the multiplex (400 bp amplicon) approach. Regions of reduced accuracy in the Nanopore-generated consensus sequence (indicated by the blue line and plotted on the right y-axis) were correlated with regions of low sequencing coverage (indicated by the area plot and plotted against the left y-axis). Consensus accuracy was assessed by alignment against a reference sequence, generated using the standard, Illumina and single-plex (2 kb amplicon), approach. For DENV1, 2 and 4, the accuracy of the 3’ end of the consensus sequences could not be fully assessed, as the single-plex approach did not generate a complete sequence.

### Phylogenetic analysis of Nanopore-and Illumina-generated consensus sequences

Phylogenies were constructed for each DENV serotype using the complete coding regions, to assess whether the consensus sequences produced by the Nanopore multiplex method and the Illumina single-plex method produced similar results. The consensus sequences generated using the two approaches formed monophyletic clusters (Figure 4). The consensus sequences generated from the control samples were more closely related (mean distance between taxa = 0.19 %, range 0.04 – 0.39 %), than those generated from the clinical samples (mean distance between taxa = 1.19 %, range 0.43 - 2.29 %).

**Figure 4.**
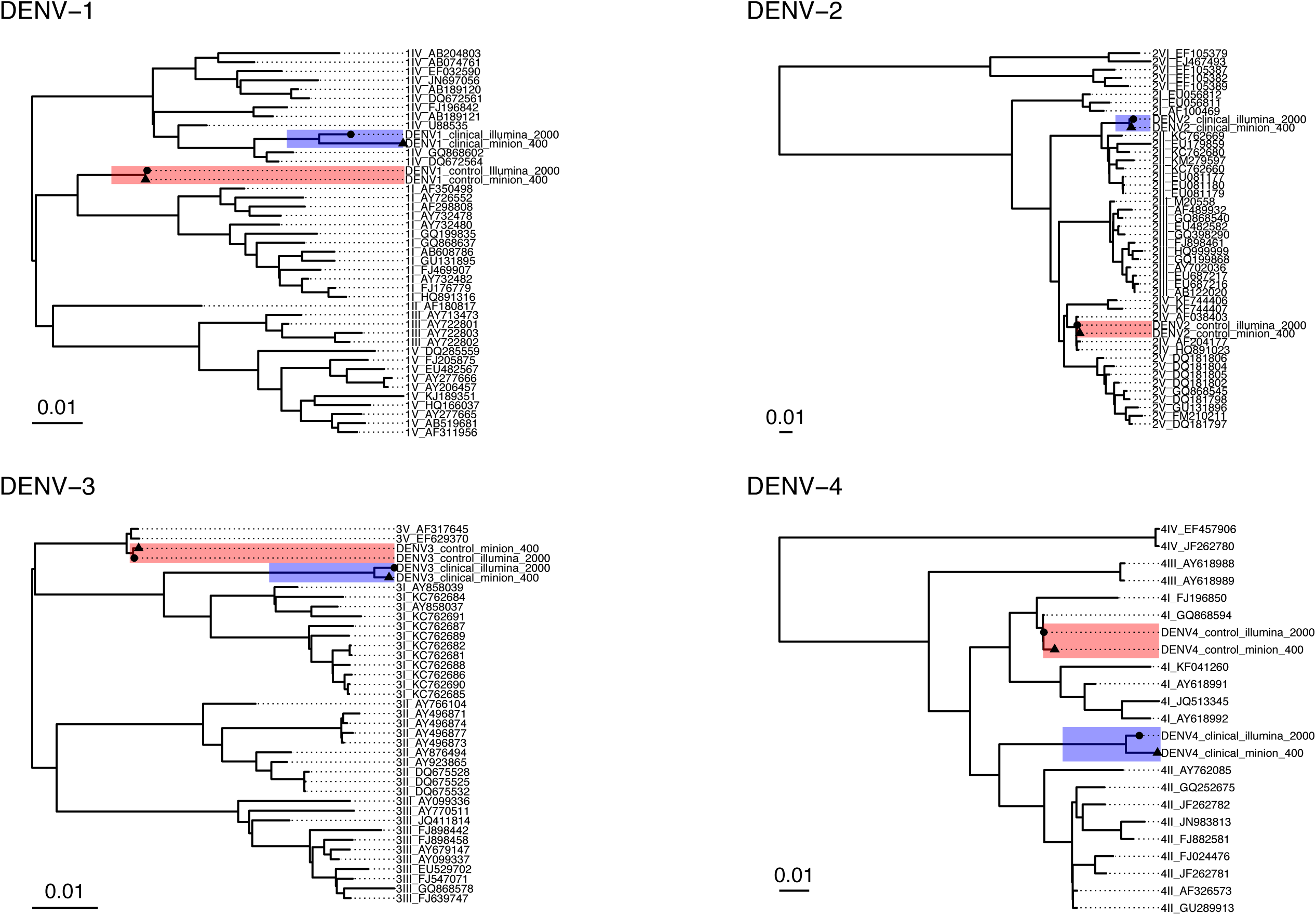
Phylogenetic analysis of DENV1-4 consensus sequences. Consensus sequences generated using the current DENV sequencing approach (Illumina-sequencing of single-plex (2 kb) amplicons) are depicted with a circle (•), and the newly developed approach (Nanopore-sequencing of multiplex (400 bp) amplicons) with a triangle (▲). Phylogenies were generated using the complete DENV coding regions. Sequences generated from control RNA samples, and those generated from real clinical samples, formed monophyletic clades (highlighted in red and blue respectively).

## Discussion

Here, we aimed to develop a fast, robust and portable approach for DENV sequencing, to be used in laboratories with limited resources, or in the field. We used the ‘Primal Scheme’ tool to design multiplex primer sets capable of amplifying the DENV coding region in two simultaneous reactions. The method was compared to the current, single-plex approach, by sequencing the resulting amplicons on both the Illumina MiSeq and Oxford Nanopore Technologies MinION platforms.

The multiplex PCR approach in combination with Nanopore sequencing was robust, generating full-genome coverage for all DENV isolates at the first attempt with no optimisation of primer concentrations or thermal cycling conditions. Conversely, the single-plex approach performed well for the control samples, but failed to produce several amplicons from the clinical isolates. In order to generate the missing amplicons, the reactions had to be repeated with alternative primers taken from the multiplex set. As demonstrated here, the single-plex approach makes it possible to validate each reaction individually, which can be useful for guaranteeing full genome coverage before sequencing begins. However, in a situation where resources are limited, or when a result is urgently required, the robustness of the multiplex approach may be advantageous.

The primer sets were designed based on DENV isolate sequences reported from Indonesia, as this is the region our future work will be focussed on. However, the primers were successfully tested on clinical samples taken from patients in the Phillippines, suggesting that the method is robust to some sequence variation, within the South-East Asian isolates at least. Using the primer sets on more genetically divergent DENVs such as isolates from South American or Africa should also be possible. However, the reactions may require some primers to be replaced in order to achieve the best results.

The multiplex approach was assessed in combination with the Nanopore MinION sequencing device as the device’s portability, low-cost and ease of use make it an excellent candidate for sequencing in the field and other resource-limited settings. A notable disadvantage of Nanopore sequencing however, is its high error rate compared to other platforms such as Illumina (30). Analysis of the Nanopore data generated for this study revealed that the individual reads were only 90 % accurate when compared to the final Illumina-generated consensus sequence. However, assembling the Nanopore reads into consensus sequences significantly improved upon the raw read accuracy, generating sequences ≥ 99.5 % identical to those produced by the Illumina single-plex method.

Interestingly, when sequenced on the Nanopore platform, the multiplex amplicon method routinely produced marginally less accurate consensus sequences compared to the single-plex approach, despite comparable raw read accuracies and genome coverage. This appeared to be due to a difference in coverage depth between the two methods: depth of coverage in the multiplex dataset was not uniform across the DENV genome, instead exhibiting distinct peaks and troughs. These regions of low coverage corresponded with regions of reduced accuracy in the consensus sequence, presumably the coverage in these regions was not enough to overcome the inaccuracies in the Nanopore reads. Conversely, coverage depth of the single-plex approach was both uniform and high, due to the ability to pool amplicons in an equimolar fashion.

It should be possible to improve coverage depth by replacing problem primers or adjusting their concentrations. However, it is also worth noting that the MinION runs in this study were stopped after 8 – 12 h, yet each flow-cell is capable of running for up to 48 hours. The runs were terminated early as the read count was deemed sufficient for reconstructing the DENV genomes, and the remaining sequencing capacity could be used for future runs as a cost saving measure. However, it may be possible to improve coverage depth further by simply allowing the MinION to run for a longer period of time. Indeed, in this scenario the ability to monitor sequencing in real-time is a major benefit of MinION sequencing compared to Illumina-based methods. To this end, it would be beneficial to use software capable of assessing coverage in real-time (e.g. mapping to a reference genome) so that sufficient coverage can be ensured before stopping the sequencing run (e.g. use of github.com/artic-network/rampart).

Based on these results, it may be important to mask regions of low coverage if a highly accurate consensus sequence is required (e.g. for calling SNPs). The accuracy of the consensus sequences may also be improved by using a tool such as Nanopolish (available at: nanopolish.readthedocs.io/en/latest/). Nanopolish is a software package designed to improve the accuracy of Nanopore-generated consensus sequences by referring back to the event-level ‘squiggle’ data in order to resolve SNPs, insertions and deletions, and has been found to work well for similar amplicon-based, viral data-sets (31). However, despite testing various iterations of the software, we were unable to improve the accuracy of our consensus sequences. Regardless, phylogenetic analysis demonstrated that any inaccuracies in the Nanopore-generated consensus sequences did not prevent them from generating monophyletic clusters with their single-plex, Illumina-generated counterparts. This is important to note as Nanopolish is somewhat computationally intensive to run and may not even be possible in settings with limited computing resources. Additional accuracy may be obtained through improvements in basecalling and read mapping; we found that the graphmap program gave more accurate consensus sequences than the more widely used BWA and minimap2 programs, at least in part due to higher sensitivity, leading to fewer regions of low coverage (i.e. below 50 x) (Supplementary Table 1). Although the permissiveness of graphmap may lead to false positives in direct sequencing studies, this is less likely to pose a problem in amplicon-based sequencing of short viral genomes.

Overall, the multiplex method was found to be a robust and efficient method for generating full-genome sequences of DENV. However, due to uneven coverage, the consensus sequences were slightly less accurate than those generated using the conventional, single-plex approach. However, it should be noted that, in our hands, the single-plex method was time consuming and unreliable for several clinical samples. Therefore, although marginally less accurate, the multiplex approach may be preferred when sequencing unknown or unusual isolates, when working in low-resource settings, or when rapid results are required.

Despite its higher error rate, Nanopore sequencing produced consensus sequences highly similar to those generated by the Illumina platform, demonstrating that, coupled with its portability and affordability, the MinION device is a viable tool for on-site virus surveillance in resource-limited settings where DENV is prevalent. However, the accuracy and multiplexing potential of the Illumina platform mean that it is likely to remain the preferred option in well-equipped central laboratories.

Finally, as the multiplex method produces smaller amplicons, it may be useful for working with partially degraded samples which cannot be amplified using the single-plex approach. The method may also be suitable for studies wanting to sequence DENV from asymptomatic individuals from whom, due to having low viral loads, it is difficult to generate sequences, yet are thought to represent the majority of DENV infections (2). These asymptomatic individuals have been shown to be capable of transmitting the virus (32), and could therefore potentially be acting as hidden reservoirs during outbreaks.

## Supporting information

Dengue multiplex primer sets

## Acknowledgements

This study was co-funded by the MRC and the Indonesian Science Fund (DiPi) as part of the UK-Indonesia Joint Health Research Call on Infectious Diseases (2016).

The authors would like to acknowledge the team at the Cambridge Stratified Medicine Core Laboratory for providing the Illumina library preparation and sequencing services.

