## Supplementary Table and Figures for "A Nanopore-based method for generating complete coding region sequences of dengue virus in resource-limited settings"

| Virus (length) | Mapping Algorithm [a] | > 5 x Coverage (%) [b] | > 50 x Coverage (%) [c] | Consensus Accuracy (%) [d] |
|---|---|---|---|---|
| DENV1 (10652 bp) | Graphmap | 100.00 | 99.41 | 99.92 |
| | BWA | 100.00 | 97.77 | 99.88 |
| | Minimap2 | 100.00 | 95.16 | 99.89 |
| DENV2 (10642 bp) | Graphmap | 100.00 | 93.14 | 99.67 |
| | BWA | 100.00 | 90.83 | 99.63 |
| | Minimap2 | 97.48 | 87.92 | 97.35 |
| DENV3 (10641 bp) | Graphmap | 100.00 | 90.25 | 99.86 |
| | BWA | 100.00 | 88.44 | 99.90 |
| | Minimap2 | 98.48 | 86.60 | 98.37 |
| DENV4 (10507 bp) | Graphmap | 100.00 | 91.61 | 99.52 |
| | BWA | 100.00 | 90.98 | 99.23 |
| | Minimap2 | 95.08 | 90.66 | 98.56 |

[a] Mapping algorithm used to generate consensus sequence

[b] Percentage of the viral genome covered by 5 or more reads. Coverage below this threshold was insufficient to call a consensus.

[c] Percentage of the viral genome covered by 50 or more reads. When coverage is below this threshold, consensus accuracy can be significantly affected

[d] Similarity to an Illumina-generated reference sequence, determined by alignment

**Supplementary Table 1. A comparison of mapping algorithms for Nanopore data derived from amplicons.** Three mapping algorithms: graphmap v 0.5.2, BWA v 0.7.17, and minimap2 v 2.9 were tested using the same sets of Nanopore reads. Reads were generated from dengue virus control RNA, using the multiplex (400 bp) approach. Graphmap and BWA mapped sufficient reads to generate a minimum of 5 x coverage depth across 100 % of the dengue virus coding region for all serotypes. Minimap2 produced the worst coverage, which negatively affected consensus accuracy. 50 x coverage was observed to be the threshold below which consensus accuracy is likely to be negatively affected. Graphmap produced the greatest 50 x or more coverage for all four DENV serotypes. Comparison of the consensus sequences to an Illumina-generated reference revealed that graphmap also produced the most accurate consensus sequence for three of the four isolates. BWA did however generate the most accurate consensus for DENV-3 (99.90 % compared to 99.86% for graphmap). Based on these results, graphmap was selected for use in this study.

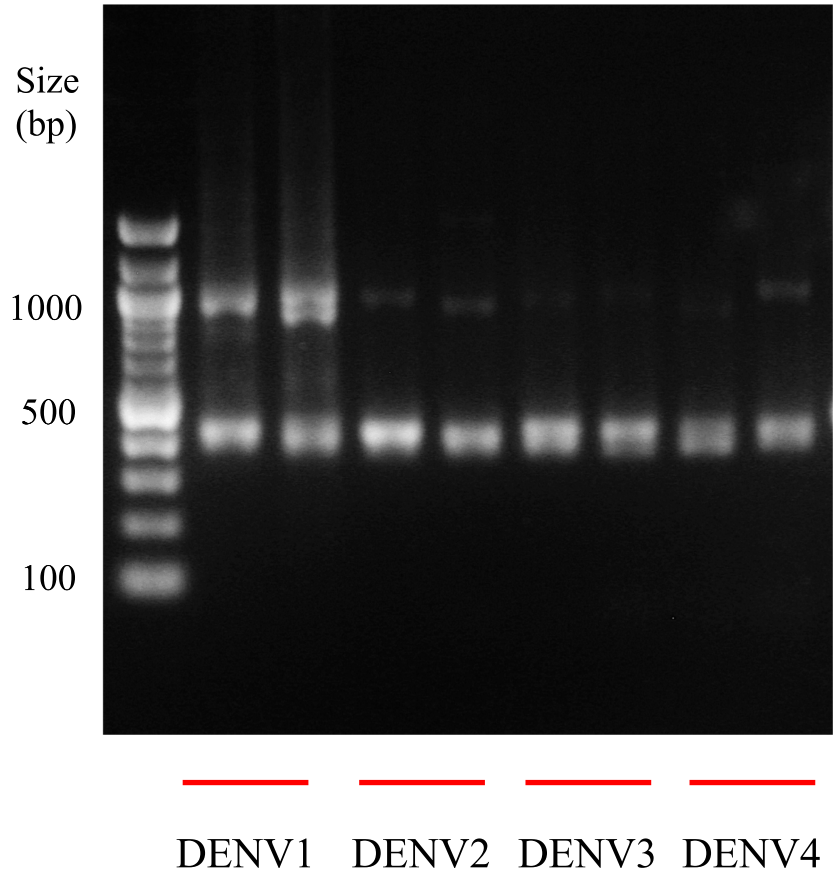


**Supplementary Figure 1. Multiplex PCR of DENV1-4 control RNA samples.** Two PCR reactions were performed for each sample to keep overlapping amplicons separate. All reactions produced the expected 400 bp band. A band approximately 1 kb in size can also be seen for DENV1, and faintly for DENV2-4, demonstrating that some larger amplicons had been generated.

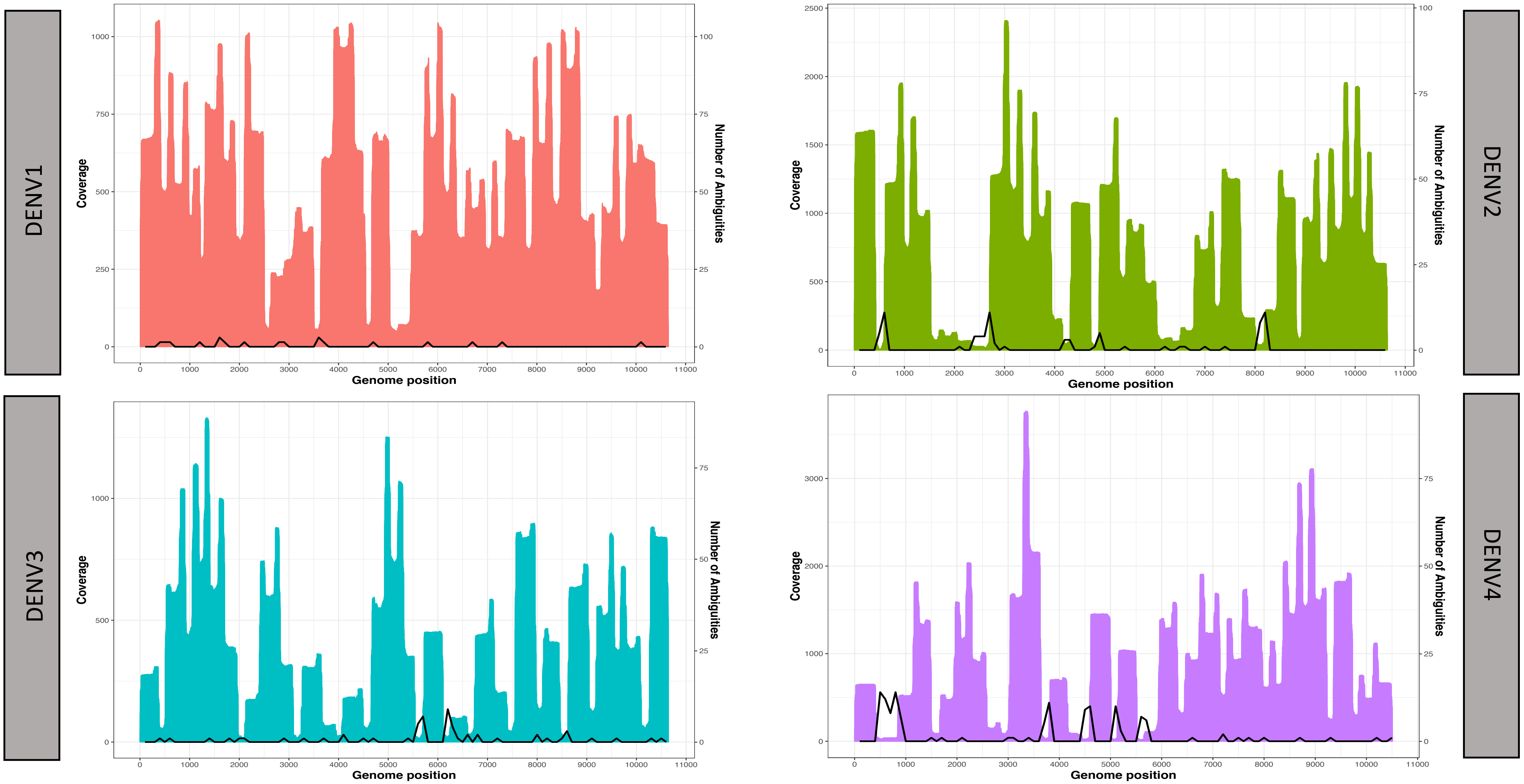


**Supplementary Figure 2. Relationship between depth of sequencing coverage and ambiguous base-calls in Nanopore-generated consensus sequences.** DENV control samples were sequenced using the multiplex PCR (400 bp amplicon) approach. Regions of the consensus sequences containing high numbers of ambiguous base-calls (indicated by the black line and plotted on the right y-axis) coincided with regions of reduced sequencing coverage (indicated by the area plot and plotted against the left y-axis).

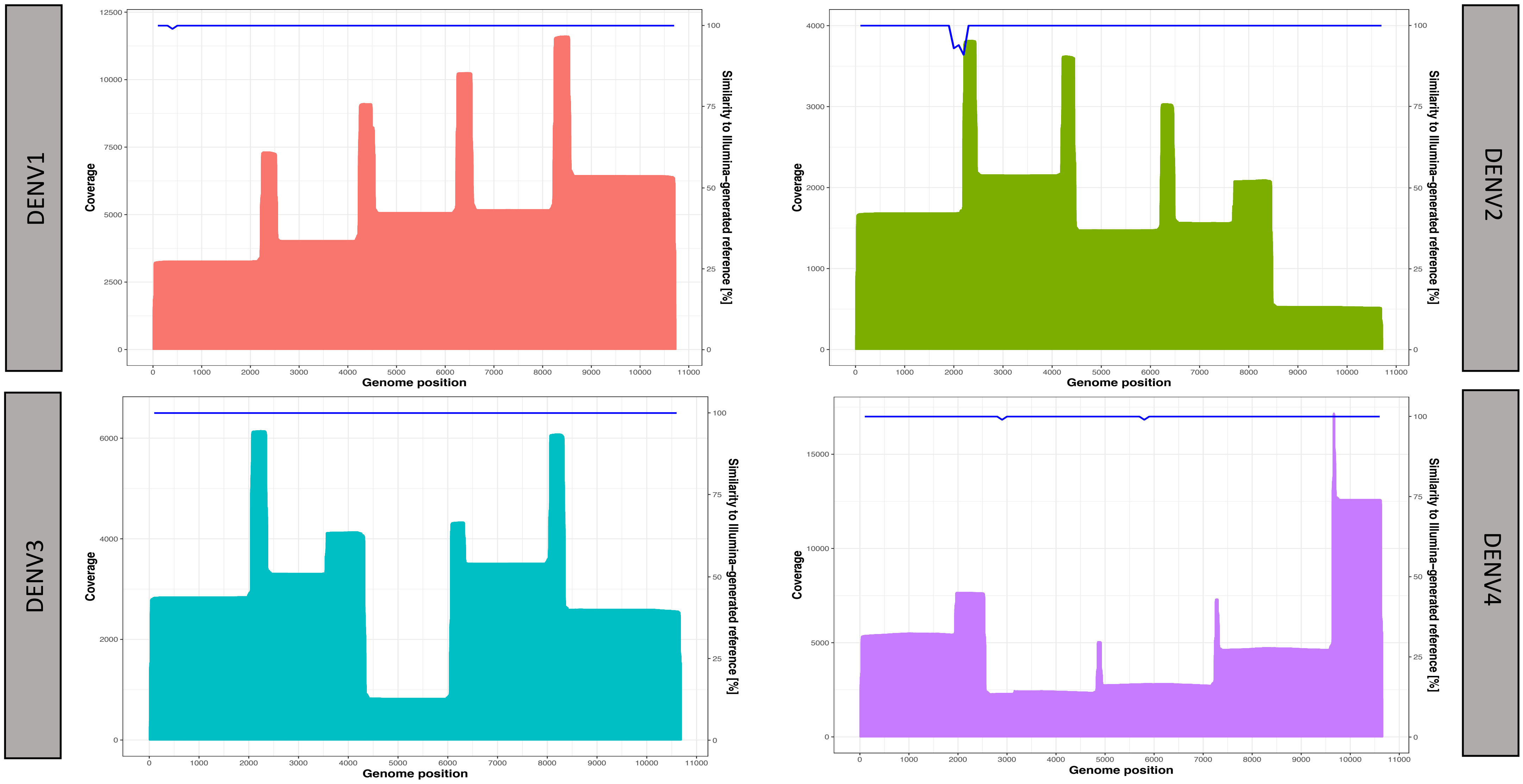


**Supplementary Figure 3. Nanopore sequencing coverage and accuracy for DENV control samples, using the single-plex PCR (2 kb amplicon) approach.** Accuracy of the consensus sequences was determined by comparison to reference sequences generated using the single-plex PCR and Illumina sequencing approach. Drops in consensus sequence accuracy were rare in the consensus sequence (indicated by the blue line and plotted on the right y-axis) as sequencing coverage was uniformly high (indicated by the area plot and plotted against the left y-axis).
