## Supplementary material for "A Nanopore-based method for generating complete coding region sequences of dengue virus in resource-limited settings": Dengue multiplex primer sets

| Serotype | Primer | Sequence |
| --- | --- | --- |
| DENV1 | DENV1_1_LEFT | AGTCTACGTGGACCGACAAGAA |
| DENV1 | DENV1_1_RIGHT | GGCATCAGCATAAGGAGCATGG |
| DENV1 | DENV1_2_LEFT | TGGCTAGATGGGGCTCATTCAA |
| DENV1 | DENV1_2_RIGHT | TCGCCAGTTTGGGAACATGTTC |
| DENV1 | DENV1_3_LEFT | GGATTTGGGAGAGTTATGTGAGGAC |
| DENV1 | DENV1_3_RIGHT | TCCTGACAGTCCTTCCACGAAG |
| DENV1 | DENV1_4_LEFT | GCACATGCCATAGGAACATCCA |
| DENV1 | DENV1_4_RIGHT | ACGTTCGTCGACACACAAAGTT |
| DENV1 | DENV1_5_LEFT | TGGACATTGAACTCTTGAAGACGG |
| DENV1 | DENV1_5_RIGHT | TGCAGTTGTCCCATGTTCTGTG |
| DENV1 | DENV1_6_LEFT | AGTGTGTGACAAAACTGGAAGGAA |
| DENV1 | DENV1_6_RIGHT | TCCTGCTTCTTTGCATGAGCTG |
| DENV1 | DENV1_7_LEFT | TGGCTAGTCCACAAACAATGGTTT |
| DENV1 | DENV1_7_RIGHT | CATGGTGCATCTGTTCCTTCGT |
| DENV1 | DENV1_8_LEFT | AGATGGACAAACTGACTCTAAAAGGG |
| DENV1 | DENV1_8_RIGHT | TCCAACAGACGTGAACACTCCT |
| DENV1 | DENV1_9_LEFT | AGCTGGTTCAAGAAAGGAAGCA |
| DENV1 | DENV1_9_RIGHT | TGCTCTGTCCAAGTGTGAACTT |
| DENV1 | DENV1_10_LEFT | TGCAGTTGGCCTAGTAACGCTA |
| DENV1 | DENV1_10_RIGHT | GTTCCATGGGTTGTGGCCTAAT |
| DENV1 | DENV1_11_LEFT | ACATCATGTGGAAGCAAATATCAAATGAA |
| DENV1 | DENV1_11_RIGHT | TATCCAGTACCCCATGTCAGCG |
| DENV1 | DENV1_12_LEFT | TGGATTTGGAATTTTCACGACAAACA |
| DENV1 | DENV1_12_RIGHT | CCGTGGTACCCTCACACAAATC |
| DENV1 | DENV1_13_LEFT | TGATAATTCCAAAGATCTATGGAGGACC |
| DENV1 | DENV1_13_RIGHT | GCAGTCCTAGTGAAAAGCTGTCC |
| DENV1 | DENV1_14_LEFT | AGGAGAAGATGGGTGCTGGTAC |
| DENV1 | DENV1_14_RIGHT | TCTCTGGATGTTAGTCTGCGGA |
| DENV1 | DENV1_15_LEFT | TATGCATCATGGTTGGAGCCAA |
| DENV1 | DENV1_15_RIGHT | TGGACAGGCATAAGGGGAAGAG |
| DENV1 | DENV1_16_LEFT | TGACTGACTTTCAATCACATCAGCT |
| DENV1 | DENV1_16_RIGHT | GCTTCTTCTTCCCAGGAGACCT |
| DENV1 | DENV1_17_LEFT | AAAATGATGTGCCGCTAGCTGG |
| DENV1 | DENV1_17_RIGHT | ACTTCTGGAGGGCTAGGTGTGT |
| DENV1 | DENV1_18_LEFT | CTCTGCTAGCAGTTTCAGGGGT |
| DENV1 | DENV1_18_RIGHT | TCTTCTCCCGTGTTCCAGGATC |
| DENV1 | DENV1_19_LEFT | ATGTCACCAGGGGAGCTGTACT |
| DENV1 | DENV1_19_RIGHT | GCCCTTCTTGTGATGCTTTAGCT |
| DENV1 | DENV1_20_LEFT | TAAACCCGGCACATCTGGATCT |
| DENV1 | DENV1_20_RIGHT | TGAAAGTGGCGTGACACATGAG |
| DENV1 | DENV1_21_LEFT | AAGGGTTGTCGCTTCCGAAATG |
| DENV1 | DENV1_21_RIGHT | ACCAAACTGTTTTGCCTGGGAA |
| Serotype | Primer | Sequence |
| DENV1 | DENV1_22_LEFT | GCGATCTTCATGACAGCCACTC |
| DENV1 | DENV1_22_RIGHT | ATGACACGCTCTGGACCATCTT |
| DENV1 | DENV1_23_LEFT | GGGACTACGTCGTCACAACAGA |
| DENV1 | DENV1_23_RIGHT | AACGTTTTCCTTGCTTCACCCC |
| DENV1 | DENV1_24_LEFT | GGACGGAAGCAAAAATGCTCCT |
| DENV1 | DENV1_24_RIGHT | TGAGACGCTTCTTCTTCCTGCT |
| DENV1 | DENV1_25_LEFT | TGGACGTGGAGATCTGGACAAA |
| DENV1 | DENV1_25_RIGHT | GGCTCTACACTGGCCATCCATA |
| DENV1 | DENV1_26_LEFT | GCTGTGTTAACTGGTGGAGTGAC |
| DENV1 | DENV1_26_RIGHT | TTGTTGTGGCCACTGCATAGAG |
| DENV1 | DENV1_27_LEFT | GGGACTGCTGGAAACCACAAAG |
| DENV1 | DENV1_27_RIGHT | CTTTTGCTTGCAGTCCAGGTCC |
| DENV1 | DENV1_28_LEFT | TAGGAGTTCCACTTCTCGCCTT |
| DENV1 | DENV1_28_RIGHT | TGTTTGCCATGGAAACCGCTAT |
| DENV1 | DENV1_29_LEFT | TCACAGATCCTCTTGATGCGGA |
| DENV1 | DENV1_29_RIGHT | TTCCTCTCCACAAACCACCTCA |
| DENV1 | DENV1_30_LEFT | GAATTCAACACCTATAAAAGGAGTGGGATT |
| DENV1 | DENV1_30_RIGHT | TTGGGTTTGGAGAGGACTCACC |
| DENV1 | DENV1_31_LEFT | ATCCCAATGGCGACCTATGGAT |
| DENV1 | DENV1_31_RIGHT | TGGTTTCCTGTGAGCCATTGTG |
| DENV1 | DENV1_32_LEFT | CTAGTGCGAAATCCACTTTCAAGAAA |
| DENV1 | DENV1_32_RIGHT | GGCTATTTGTGTGACCATGGGG |
| DENV1 | DENV1_33_LEFT | AAAACATGGGCCTATCATGGATCA |
| DENV1 | DENV1_33_RIGHT | CTCTGTGCACAAGGTCCCAGAA |
| DENV1 | DENV1_34_LEFT | AGAGAGGAGTTTACAAGAAAAGTTAGGT |
| DENV1 | DENV1_34_RIGHT | ATCCGGCTGTGTCATCTGCATA |
| DENV1 | DENV1_35_LEFT | GACCACTGGTTCAGCAGAGAGA |
| DENV1 | DENV1_35_RIGHT | GGGCTTCCATGTTGGTGAAAGT |
| DENV1 | DENV1_36_LEFT | GGTAAGGGTACAGAGACCAGCA |
| DENV1 | DENV1_36_RIGHT | TCCATCCTTTTGAGGGTTCCCA |
| DENV1 | DENV1_37_LEFT | GCAATCAGTGGAGATGACTGCG |
| DENV1 | DENV1_37_RIGHT | CCAATCAACTGGAACGGCTGAA |
| DENV1 | DENV1_38_LEFT | GCTGGATGGAGCCTGAGAGAAA |
| DENV1 | DENV1_38_RIGHT | CACTTGTATGTTGGTGGCCCAG |
| DENV1 | DENV1_39_LEFT | CATGGATGGAGGACAAAACCCA |
| DENV1 | DENV1_39_RIGHT | TCGGCCTGACTTCATTTTACGTC |
| DENV1 | DENV1_40_LEFT | TGAGTCAACACACTTACAAAATGAAGGA |
| DENV1 | DENV1_40_RIGHT | CTGGTCTCTCCCAGCGTCAATA |
| DENV2 | DENV2_1_LEFT | TACGTGGACCGACAAAGACAGA |
| DENV2 | DENV2_1_RIGHT | AACGCCATCACTGTTGGAATCA |
| DENV2 | DENV2_2_LEFT | AAAAATCAAAGGCTATCAATGTCTTGAGA |
| DENV2 | DENV2_2_RIGHT | GGAACGAGTGCCACTGATCTTTT |
| Serotype | Primer | Sequence |
| DENV2 | DENV2_3_LEFT | CAGGCAGAATGAACCAGAAGACA |
| DENV2 | DENV2_3_RIGHT | TCAACCCAGCTTCCTCCTGAAA |
| DENV2 | DENV2_4_LEFT | GGAACGACATATTTCCAAAGAGTCCT |
| DENV2 | DENV2_4_RIGHT | AATCCGCATCCATTTCCCCATC |
| DENV2 | DENV2_5_LEFT | GAGGCAAAGCTGACCAACACAA |
| DENV2 | DENV2_5_RIGHT | TCCATTTGCAGCAACACCATCT |
| DENV2 | DENV2_6_LEFT | ACGGCAAGGAAATTAAAGTAACACCA |
| DENV2 | DENV2_6_RIGHT | GTTTGTCCATTCTCAGCCTGCA |
| DENV2 | DENV2_7_LEFT | GGATGTTGTCGTTTTAGGATCCCA |
| DENV2 | DENV2_7_RIGHT | GATGTAGCTGTCCCCGAATGGA |
| DENV2 | DENV2_8_LEFT | TGCAAGATCCCTTTTGAAATAATGGATTT |
| DENV2 | DENV2_8_RIGHT | GTGAGGTGCTGCGTGAATTCAT |
| DENV2 | DENV2_9_LEFT | CATCCATAGGAAAGGCCCTCCA |
| DENV2 | DENV2_9_RIGHT | ACTGAGCGGATTCCACAAATGC |
| DENV2 | DENV2_10_LEFT | TGTGGCAGTGGGATTTTTATCACA |
| DENV2 | DENV2_10_RIGHT | CTGTTTCGGGGCCATCAATGAG |
| DENV2 | DENV2_11_LEFT | TCATGCAGGCAGGAAAACGATC |
| DENV2 | DENV2_11_RIGHT | AGAGTGTGTGACTTTGGCCAGT |
| DENV2 | DENV2_12_LEFT | AAAAGACAACAGAGCCGTCCAC |
| DENV2 | DENV2_12_RIGHT | GTGGTAATGTGCAAGATCGGCA |
| DENV2 | DENV2_13_LEFT | TTTCTGCGAAGGAACCACAGTG |
| DENV2 | DENV2_13_RIGHT | TGTCATCCGTCATAGTAGCGCC |
| DENV2 | DENV2_14_LEFT | CCCGAGTAGGAACGAAACATGC |
| DENV2 | DENV2_14_RIGHT | CTGCATTTGGGACGCACAAGAT |
| DENV2 | DENV2_15_LEFT | AGAGCACCATACCAGAGACTATACT |
| DENV2 | DENV2_15_RIGHT | TCACCATCCCGACTGCCATAAT |
| DENV2 | DENV2_16_LEFT | TGGATACCACTGGCATTGACGA |
| DENV2 | DENV2_16_RIGHT | TCCTGATACCACCAACAGTCCTG |
| DENV2 | DENV2_17_LEFT | ACAGGCAGAGATATCAGGAAGTAGTC |
| DENV2 | DENV2_17_RIGHT | GTCCGCCCATGATGGTTCAATT |
| DENV2 | DENV2_18_LEFT | AGAAAGGGATTCTAGGATACTCGCA |
| DENV2 | DENV2_18_RIGHT | TCAGTCTGGGCTATAGCACTCAC |
| DENV2 | DENV2_19_LEFT | GCCGTGTCTCTGGACTTTTCTC |
| DENV2 | DENV2_19_RIGHT | TTAGATCCACAATCTCCCGCCC |
| DENV2 | DENV2_20_LEFT | TAATCCTGGCCCCCACTAGAGT |
| DENV2 | DENV2_20_RIGHT | CATGTCCAGAGTTCCACGAACG |
| DENV2 | DENV2_21_LEFT | TAGAGATGGGTGAAGCAGCTGG |
| DENV2 | DENV2_21_RIGHT | TTTCATACAGCGTCTGGGGTCT |
| DENV2 | DENV2_22_LEFT | CTCAGCAGGAAGACTTTTGATTCTGA |
| DENV2 | DENV2_22_RIGHT | TGGTTCGAACATGCTGGGAATG |
| DENV2 | DENV2_23_LEFT | ACCAGTACATATACATGGGAGAACCT |
| DENV2 | DENV2_23_RIGHT | CCTTGAATTCTTTGAGCGCCAG |
| Serotype | Primer | Sequence |
| DENV2 | DENV2_24_LEFT | TGGTGCTTTGATGGAGTCAAGAA |
| DENV2 | DENV2_24_RIGHT | ACATTCCCAGGGTCATCTTCCC |
| DENV2 | DENV2_25_LEFT | TAGTGAACTGCCGGAGACTCTG |
| DENV2 | DENV2_25_RIGHT | TCCAGGATGTTGCTCTCAGGTT |
| DENV2 | DENV2_26_LEFT | TAGCCATCCTTACAGTGGTGGC |
| DENV2 | DENV2_26_RIGHT | CTTGGAGTCCTGGCCCTATGAT |
| DENV2 | DENV2_27_LEFT | ATGGACATCGGAGTTCCCCTTC |
| DENV2 | DENV2_27_RIGHT | CCTGGATTTCCTTCCCACAGTG |
| DENV2 | DENV2_28_LEFT | TGGGACAAGTAATGCTCTTAGTCCT |
| DENV2 | DENV2_28_RIGHT | CAGCGTGATGGTCTGTTTCTCC |
| DENV2 | DENV2_29_LEFT | TGGAGAGAAATGGAAAAACCGGT |
| DENV2 | DENV2_29_RIGHT | TTGGTGACGACTCCCCTATGTC |
| DENV2 | DENV2_30_LEFT | CAGGACACGAAGAACCCATTCC |
| DENV2 | DENV2_30_RIGHT | GTGGCCTTCTTGTGTCTCATTGT |
| DENV2 | DENV2_31_LEFT | ACTCTCACGAAATTCCACACACG |
| DENV2 | DENV2_31_RIGHT | TGTCACCATAGGGATGACGTCC |
| DENV2 | DENV2_32_LEFT | CCAAGACCACCCATACAAAACGT |
| DENV2 | DENV2_32_RIGHT | GTGCCGACTTCCACTTGTTCTC |
| DENV2 | DENV2_33_LEFT | TGGCTCTGGAAAGAACTAGGAAAGA |
| DENV2 | DENV2_33_RIGHT | CCTTCTCCTTCCACTCCACTCA |
| DENV2 | DENV2_34_LEFT | AGAAGAAGCTAGGGGAGTTCGG |
| DENV2 | DENV2_34_RIGHT | CACGCACCACCTTGTTTTGGTA |
| DENV2 | DENV2_35_LEFT | TGGGACACAAGAATCACACTGG |
| DENV2 | DENV2_35_RIGHT | GCGCTTGCAAACCTGTCATCTA |
| DENV2 | DENV2_36_LEFT | TCCAGCAACTGACAGTCACAGA |
| DENV2 | DENV2_36_RIGHT | AGGTCACGTCTGTGGAAGTACA |
| DENV2 | DENV2_37_LEFT | GATGAACTGATTGGTAGGGCCC |
| DENV2 | DENV2_37_RIGHT | CAATCAATGAGCCGCACCATTG |
| DENV2 | DENV2_38_LEFT | GACATGCTGACAGTCTGGAACA |
| DENV2 | DENV2_38_RIGHT | GGGGCTCACAGGTAGCATAGTT |
| DENV2 | DENV2_39_LEFT | AAGATTTAGAAGAGAAGAGGAAGAGGCA |
| DENV2 | DENV2_39_RIGHT | GCAGGATCTCTGGTCTTTCCCA |
| DENV3 | DENV3_1_LEFT | TACGTGGACCGACAAGAACAGT |
| DENV3 | DENV3_1_RIGHT | AGCGATGTCTTTTTCCGTCTGT |
| DENV3 | DENV3_2_LEFT | ACAGCGGGAATCTTGGCTAGAT |
| DENV3 | DENV3_2_RIGHT | CACCAGCAGTCAATGTCTTCAGG |
| DENV3 | DENV3_3_LEFT | GCCTCTGGAATCAACATGTGCA |
| DENV3 | DENV3_3_RIGHT | CTCATTGTCATGGATGGGGTGAC |
| DENV3 | DENV3_4_LEFT | AGGTAGAGACATGGGCCTTTAGG |
| DENV3 | DENV3_4_RIGHT | GTAGTTCTGGTCCTGCTCCTCA |
| DENV3 | DENV3_5_LEFT | ATAGAGCTCCAGAAGACCGAGG |
| DENV3 | DENV3_5_RIGHT | TTATCTCAGCCGTGACTCCCTG |
| Serotype | Primer | Sequence |
| DENV3 | DENV3_6_LEFT | TGTTTGGAATTAATAGAGGGAAAAGTGGT |
| DENV3 | DENV3_6_RIGHT | CTGTGTGCATTGCTCCCTCTTG |
| DENV3 | DENV3_7_LEFT | ATGGACATCAGGGGCTACAACA |
| DENV3 | DENV3_7_RIGHT | GCTGTGATCAGTCTGCCATTGT |
| DENV3 | DENV3_8_LEFT | AGTCTCTGAAACACAACATGGGAC |
| DENV3 | DENV3_8_RIGHT | GGCCGTGTAAGCACTTCCAAAT |
| DENV3 | DENV3_9_LEFT | CGATTGGGAAGATGTTCGAGGC |
| DENV3 | DENV3_9_RIGHT | TCTTTTGGGGGAGTCTGCTTGA |
| DENV3 | DENV3_10_LEFT | ACATGGGGTGTGTCATAAACTGG |
| DENV3 | DENV3_10_RIGHT | CTGGTGTGTTTGGCCCATCTAT |
| DENV3 | DENV3_11_LEFT | AAGAACACTAACACCACAACCCA |
| DENV3 | DENV3_11_RIGHT | GTGTGTGATTTTGGCCATGTGC |
| DENV3 | DENV3_12_LEFT | AAGGATGAGAGAGCCGTACACG |
| DENV3 | DENV3_12_RIGHT | TATCGCAGGGGAGGAAGTGTAC |
| DENV3 | DENV3_13_LEFT | AGGAACAACAGTTGTCATCACAGA |
| DENV3 | DENV3_13_RIGHT | CATTCCCATTCTGTCAGAGGCG |
| DENV3 | DENV3_14_LEFT | AAAAGCACATGATTGCAGGGGT |
| DENV3 | DENV3_14_RIGHT | GTTCTCCAGGCAACAGTCAACG |
| DENV3 | DENV3_15_LEFT | ATGGCGAATGGAATAGCCTTGG |
| DENV3 | DENV3_15_RIGHT | AGCCATGGGTACGTCATTCCTAA |
| DENV3 | DENV3_16_LEFT | TCCACCTCTACCACTTTTTATTTTCAGT |
| DENV3 | DENV3_16_RIGHT | GGGTTTGCTTTTGCCAAGTGTG |
| DENV3 | DENV3_17_LEFT | TGAGAATAAAAGATGATGAGACTGAGAACA |
| DENV3 | DENV3_17_RIGHT | GCACCTCTTCTCCTTTTTGCCA |
| DENV3 | DENV3_18_LEFT | CGTCACAAGAGGAGCAGTGTTG |
| DENV3 | DENV3_18_RIGHT | TTCTTCCAACTCTGGTGTCGGT |
| DENV3 | DENV3_19_LEFT | ACAGAGAGGGAAAGGTAGTGGG |
| DENV3 | DENV3_19_RIGHT | GACAGCAAACGCATTGTGAACG |
| DENV3 | DENV3_20 | TTGCAGCTGAGATGGAAGAAGC |
| DENV3 | DENV3_21 | GAGAGGCAGCCGCAATTTTCAT |
| DENV3 | DENV3_22 | ACTAAACTAAATGATTGGGACTTTGTGGT |
| DENV3 | DENV3_23 | TGCTGCTAGACAACATCAACACA |
| DENV3 | DENV3_24 | TGGATGTGGAAATCTGGACAAAGG |
| DENV3 | DENV3_25 | GGCATGCAGTGGAGGAATTACC |
| DENV3 | DENV3_26 | AGCAGAGAACTCCCCAAGACAA |
| DENV3 | DENV3_27 | GCAGTGGTCCTGATGGGTTTAG |
| DENV3 | DENV3_28 | AGACCTAGATCCTGTAATATACGACTCA |
| DENV3 | DENV3_29 | AGGAACAGGGTCACAAGGTGAA |
| DENV3 | DENV3_30 | GTGCGAGGATACACAAAAGGCG |
| DENV3 | DENV3_31 | TACATGCCAGCTGTGATTGAGC |
| DENV3 | DENV3_32 | TCAATGCGGAACCAGAAACACC |
| DENV3 | DENV3_33 | AGAGAAAGTGGACACCAGGACA |
| Serotype | Primer | Sequence |
| DENV3 | DENV3_34 | GGGCAAGAGAGAGAAGAAACTTGG |
| DENV3 | DENV3_35 | AGAAGATGACCTGCACAATGAGG |
| DENV3 | DENV3_36 | TCGAAGACAGACCTCGAGAACC |
| DENV3 | DENV3_37 | CCTTTCTGCTCCCACCACTTTC |
| DENV3 | DENV3_38 | ACAGAAGACATGCTTACTGTTTGGAA |
| DENV3 | DENV3_39 | AAAAGAGGCAAACTGTCAGGCC |
| DENV4 | DENV4_1 | TGTTAGTCTGTGTGGACCGACA |
| DENV4 | DENV4_2 | GGGACAGTTGAAAAAGAATAAGGCCA |
| DENV4 | DENV4_3 | CCTGGGTGAAATGTGTGAAGACA |
| DENV4 | DENV4_4 | GCTCCTGGCAGGATTTATGGCT |
| DENV4 | DENV4_5 | TAACCACGGCAACAAGATGTCC |
| DENV4 | DENV4_6 | GGAGACACCCATGCAGTAGGAA |
| DENV4 | DENV4_7 | TTCCTCATGCCAAGAGACAGGA |
| DENV4 | DENV4_8 | AGTTCCCATAGAGATAAGAGATGTGAACA |
| DENV4 | DENV4_9 | GGCCATTCTAGGTGAAACAGCT |
| DENV4 | DENV4_10 | GTGGAAGCGGAATCTTTGTGGT |
| DENV4 | DENV4_11 | ACCTCCAGTGAATGATCTGAAATATTCAT |
| DENV4 | DENV4_12 | ACATGGGTTATTGGATAGAGAGCTCA |
| DENV4 | DENV4_13 | AGGATTGTGACCATAGAGGCCC |
| DENV4 | DENV4_14 | ACCCTGTTCGTGGAAGAATGCT |
| DENV4 | DENV4_15 | AATGGCCATGACAACGGTGTTT |
| DENV4 | DENV4_16 | GTCCCATTGGGTAGAAATAACAGCA |
| DENV4 | DENV4_17 | CAATGGGATGAAATGGCGGACA |
| DENV4 | DENV4_18 | AGGATCATGCAAAGAGGGTTGTT |
| DENV4 | DENV4_19 | GGAGAAATTGGAGCAGTAACTCTGG |
| DENV4 | DENV4_20 | TCTCCCATCAATAGTCAGAGAAGCT |
| DENV4 | DENV4_21 | CCCTTGTAGTGTCGCAGCTAGA |
| DENV4 | DENV4_22 | AGTCGGGAAAGAGAGTGATCCAG |
| DENV4 | DENV4_23 | ATGTTTTCTCCGGAGACCCACT |
| DENV4 | DENV4_24 | AAGACCGAGAATGGTGCTTCAC |
| DENV4 | DENV4_25 | AGCCCTTGACAACATAGTCATGC |
| DENV4 | DENV4_26 | GTGCTGTTGATACCAGAACCAGAA |
| DENV4 | DENV4_27 | CCAACCAAGCAGCTGTCCTAAT |
| DENV4 | DENV4_28 | CGGTGGACGGGATAACAGTAATAGA |
| DENV4 | DENV4_29 | AACTGGGACCACAGGAGAGACA |
| DENV4 | DENV4_30 | ACACTCAAGAACGTGACTGAAGTG |
| DENV4 | DENV4_31 | AGTCCTCAATCCCTACATGCCG |
| DENV4 | DENV4_32 | GTGTCTCCACTGAAACAGAAAAACC |
| DENV4 | DENV4_33 | GTGGATACCAGAACACCACAACC |
| DENV4 | DENV4_34 | TGACAAAGAAAGGGCTCTGCAC |
| DENV4 | DENV4_35 | TCTTGGAGGACATAGACAAGAAGGA |
| DENV4 | DENV4_36 | AAGAGGTAGTGGACAGGTTGGA |
| Serotype | Primer | Sequence |
| DENV4 | DENV4_37 | CTTGAATGATATGGGAAAGGTGAGGA |
| DENV4 | DENV4_38 | GCCCAGATGTGGTCGCTTATGT |
| DENV4 | DENV4_39 | GGACTTTCTTCCAGAGCCACCT |
